# Anti-Biofouling Albumin Amyloid Coatings Enable Vancomycin-Mediated Bacterial Eradication on Medical Tubing

**DOI:** 10.1101/2025.09.14.676079

**Authors:** Ivon Y. Calibio Giraldo, Fiorela Ghilini, Eduardo Prieto, Carolina Díaz, Patricia L. Schilardi

## Abstract

Indwelling medical devices such as catheters and endotracheal tubes are major drivers of hospital-acquired morbidity and mortality due to bacterial colonization. The resulting healthcare-associated infections (HAIs) are further exacerbated by rising antimicrobial resistance, underscoring the urgent need for strategies that both prevent biofilm formation and reduce reliance on antibiotics. Polyvinyl chloride (PVC), a widely used material in medical tubing, is highly prone to bacterial attachment, making it a critical target for intervention. Here, we show that complete eradication of *Staphylococcus aureus*—both sessile on PVC and planktonic—can be achieved using a bovine serum albumin (BSA) amyloid-like coating in combination with vancomycin at a dose eight times lower than the MIC. While the amyloid film alone markedly reduces bacterial adhesion, the residual bacterial load still reaches infection-risk thresholds. This dual approach therefore not only prevents biofilm development but also significantly lowers antibiotic requirements, reducing the risk of resistance emergence and improving therapeutic safety. The coatings, deposited on PVC and on glass (as a model surface), were synthesized using dithiothreitol (DTT) as a reducing agent, as alternative to tris(2-carboxyethyl)phosphine (TCEP). Through optimization of the synthesis, the resulting films preserved their physicochemical and anti-biofouling properties while offering a simple, low-cost, and scalable approach. The coatings strongly adhere to both substrates, remain stable under aqueous and mechanical stress, and effectively suppress bacterial and mammalian cell adhesion without cytotoxicity. These properties are clinically relevant, reducing infection risk and mitigating tubing failure due to fibrous capsule formation, encasement, or crystallized biofilm-induced blockage. The demonstrated biocompatibility, robustness, and scalability of this coating platform underscore its translational potential as a clinically relevant strategy to mitigate HAIs, extend the functional lifetime of medical tubing, and alleviate the global burden of antimicrobial resistance.

## 1. Introduction

Infections associated with indwelling medical devices bring significant challenges in healthcare settings, leading to increased morbidity, mortality, and medical management costs. According to the 2020 Report from the Centre for Disease Control and Prevention (CDC) ^1^ about 50 to 70% of healthcare-associated infections (HAIs) are attributed to medical device-related infections. In hospitals alone, 1 in 31 hospitalized patients have at least one HAI, with more than 680000 infections and billions of dollars in excess healthcare costs related to HAIs treatment ^1,2^. These HAIs include central-line associated bloodstream infections, catheter-associated urinary tract infections, and ventilator-associated pneumonia. Infections may also occur at surgery sites, known as surgical site infections. The problem of device-related infections arises from the ability of bacteria to adhere to surfaces and form biofilms. Urinary and intravascular catheters are two of the most used invasive medical devices and microbial colonization of their surface is responsible for most of the HAIs ^3^. It was found that in intensive care units, urinary and bloodstream infections are catheter-associated in 96.7% and 43.3% ^4^. Moreover, lower respiratory tract infections account for 15% of all HAIs ^1^. Patients requiring mechanical ventilation are at risk of developing ventilator-associated pneumonia (VAP), largely due to the use of endotracheal tubes (ETTs). During the COVID-19 pandemic, approximately 45% of patients infected with the SARS-CoV-2 virus developed VAP ^5^. The rapid colonization of catheters’ surface and subsequent biofilm formation play a crucial role in the pathogenesis of infections and its frequent relapses. In catheterization, the biofilm may originate from the organisms that colonize the skin at the point of catheter insertion or their migration through or around the device after implantation. Biofilms are complex microbial communities encased in a protective extracellular matrix, making them highly resistant to traditional antibiotic treatments. The indiscriminate use of antibiotics has contributed to the emergence of antibiotic-resistant strains, significantly reducing the efficacy of conventional therapies. Consequently, there is an urgent need for alternative strategies that can prevent bacterial adhesion and enhance the effectiveness of antibiotics in combating biofilm formation. The use of antimicrobial coatings on medical device surfaces has emerged as a promising solution to address these challenges. These coatings offer a multifaceted approach to address the threat associated with device-related infections ^6–9^. Firstly, they could act as physical barriers, preventing bacterial adhesion and subsequent biofilm formation. Secondly, they may also hold inherent antimicrobial properties, directly targeting and eliminating bacteria that come in contact with the coating. By combining these dual functionalities, antimicrobial coatings have the potential to significantly reduce the risk of infection. An ideal coating should present several key characteristics, including antimicrobial efficacy, biocompatibility, low cost, and mechanical and long-term stability ^10^.

In recent years, self-assembled amyloid structures have emerged as promising candidates for developing innovative materials with specialized functions in a broad and diverse universe of applications ^11–14^. Amyloid structures are typically associated with neurodegenerative diseases, such as Alzheimer’s and Parkinson’s disease. However, researchers have discovered that not only the typical toxic amyloidogenic proteins but also non-toxic proteins can form amyloid-like structures with remarkable and unique properties ^15,16^. Proteins such as beta-lactoglobulin, lysozyme, and albumin can form amyloid-like fibrils that exhibit mechanical stability, biodegradability, and the ability to self-assemble into thin coatings ^17,18^.

Albumin is the most abundant protein in plasma and plays important metabolic roles. Albumin is a very stable and highly soluble protein tolerant of high temperatures. Thanks to its disulphide bonds and sulfhydryl groups, albumin can be described as chemically attractive, allowing to exploit specific chemical reactions. Albumin is one of the most studied protein, being employed in various biotechnological applications^4,19,20^. The self-assembly properties of amyloid-like albumin fibrils offer several advantages for coating applications. The process of self-assembly is pH-dependent and can be precisely controlled by varying the pH and concentration of the amyloid solution^20^. This enables the fabrication of coatings with tailored properties, including surface roughness and charge, which play crucial roles in preventing bacterial adhesion. Additionally, it has been shown that amyloid coatings exhibit substrate independence, resistance to harsh conditions, and programmable functionalities.

A coating prepared by combining amyloid-like albumin fibrils on surfaces as coadjuvant of conventional antimicrobial treatment could offer a synergistic effect, resulting in a coating with anti-adhesive properties that enhances the antibiotic effect. This work aims to introduce and explore the potential of the anti-biofouling amyloid coating on polyvinyl chloride (PVC) for biomedical applications. PVC continues to be a leading material in healthcare due to its exceptional properties that align with the industry’s strict performance requirements while remaining budget-friendly ^21,22^. Although researchers are exploring alternative materials, PVC’s proven safety and adaptability make it a reliable choice for numerous medical applications. We present the fabrication of amyloid like albumin aggregates self-assembled into a coating following a straightforward disulfide reaction step at 37°C. Previous studies have employed tris(2-carboxyethyl)phosphine (TCEP) to generate antifouling BSA-amyloid films on surfaces ^23^; here, we optimized the synthesis by using dithiothreitol (DTT) as a reducing agent. Importantly, this alternative route preserves the physicochemical, and antibiofouling properties of the resulting coatings. Furthermore, when PVC surfaces modified with BSA amyloid films and colonized by bacteria were subsequently treated with vancomycin, complete bacterial eradication was achieved. We developed a facile, biocompatible and inexpensive method to prepare an antimicrobial coating with anti-adhesive characteristics able to enhance the conventional antibiotic treatment. Consequently, this offers a comprehensive approach to prevent bacterial colonization and mitigate the risk of biofilm-associated infections. The outcomes of this research could have significant implications for the development of advanced biomaterials with broad applications in healthcare settings.

## 2 Experimental

### 2.1 BSA solution and BSA aggregates

The BSA solution was prepared by dissolving the proper amount of the protein in phosphate buffer (PB) at pH 7.02 to obtain a 40 µM BSA solution. 5.0 mL of this solution, previously stored at 4°C for 40 min, was mixed with DTT until reaching a concentration of 10 mM. The reaction mixture was stored at 4 °C for 40 min and then incubated at 37°C for 6 h. From now, the resulting dispersion will be denoted as BSA+DTT

### 2.2 Glass and PVC coating

Polyvinylchloride (PVC) pieces (5 mm diameter and 3 mm thick and 10 mm long, 25 mm wide and 3 mm thick for microbiological and stability assays, respectively) were cleaned by immersion in 96 % ethanol aqueous solution for 24 h and then sonicated in ultrapure water (Milli-Q®) for ten minutes. Glass samples obtained from microscope slides 15 mm long, 5 mm wide and 1 mm thick, and 10 mm long, 25 mm wide and 1 mm thick (Carl Roth) were used for fluorescence intensity and for stability assays, respectively. The glass samples were cleaned by immersion in piranha solution (mixture of ratio of 1:3 of 30 % v/v hydrogen peroxide, Anedra, Argentina and 91 % sulfuric acid, Anedra, Argentina) for 30 minutes. Subsequently the glasses pieces were sonicated in Milli-Q® water for 30 minutes to eliminate excess of piranha and then dried at 140 °C during 1 hour. After cleaning, the substrates were immersed in the reaction mixture previously stored at 4 °C for 40 min (see section 2.1) and then incubated for 24 h at 37 C. After that, the samples were removed from the reaction solution and stored at 4 C for at least 24 h.

### 2.3 UV-vis spectroscopy and Fluorescence spectroscopy

UV-vis spectra were acquired by a PerkinElmer Lambda 35 UV-vis spectrophotometer. UV-vis signal vs time at 650 nm were recorded for the BSA and BSA+DTT dispersions. The fluorescent-specific reagent thioflavin T (ThT) was used as a probe. 10 µL ThT, 1.04 µM, Sigma Aldrich, St. Louis, MO, USA in 96% ethanol and PBS 1:3 was added to the BSA+DTT stock solution. The fluorescence intensity measurements were carried out at 37°C on a Cary Eclipse Fluorescence Spectrophotometer, with an excitation wavelength at 440 nm and emission wavelength at 485 nm, using a 5 nm slit 5 nm width for both, excitation and emission. BSA +DTT solution (without ThT) was also measured as control.

### 2.4 Transmission Electron Microscopy (TEM)

TEM imaging of the amyloid aggregates, prepared as described in section 2.1, was performed using a Zeiss EM109T microscope equipped with a Gatan ES1000W digital camera. For sample preparation, the dispersion was kept at approximately 4 °C in a Petri dish placed over an ice-filled container. The sample was diluted 1:1 with 2% glutaraldehyde in 0.1 M phosphate buffer. A 10 µL aliquot of the diluted dispersion was placed on clean Parafilm inside a sterile Petri dish. A 200-mesh copper grid coated with an ultrathin (90–100 nm) LR White resin film was floated on the droplet for 10 minutes. The grid was then washed three times with Milli-Q water (1 minute each). For contrast enhancement, the sample was stained with 2% aqueous uranyl acetate (pH 7) for 3 minutes. Excess liquid was removed using clean filter paper, and the sample was air-dried on filter paper for 15 minutes in a laminar flow hood prior to TEM observation.

### 2.5 Attenuated Total Reflectance-Fourier Transform Infrared spectroscopy (ATR-FTIR)

ATR-FTIR was performed on a Shimadzu IRTracer-100 FTIR spectrometer equipped with a MCT detector. 5 μL of a BSA solution prepared in deuterated PBS was pipetted onto the surface of the prism and the spectral recording was initiated. The BSA-DTT solution was also prepared in deuterated PBS, following the same preparation procedure performed during the coating experiments. The solutions prepared with deuterated PBS were also used to coat the PVC samples for ATR-FTIR measurements. The coating procedure was the same as previously described. To measure, the samples were inverted onto the surface of the prism and tightened with the screw. The spectra were obtained using the following parameters: resolution 2 cm^−1^; range from 1800 to 1400 cm^−1^; 100–512 scans recorded, scanning velocity: 20 kHz, zero-filling factor: 4, apodization function: a Blackman-Harris 3-term. As a reference, D_2_O cell was used.

### 2.6 Contact angle measurements

The wetting properties of the surfaces were analyzed by measuring the water contact angle in a Ramé Hart model 290 U1 series goniometer. 3 µL of Milli-Q water were poured on the glass and PVC samples at room temperature and then the static contact angle was measured.

### 2.7 Atomic Force Microscopy (AFM)

AFM imaging was carried out in air with a Nanoscope V microscope from Bruker, operating in Tapping® mode. Images were taken at a scanning rate of 1 Hz with etched silicon tips (RTESP, 8 nm nominal radius tip, 300 kHz resonance frequency and 40 N/m force constant). To estimate the thickness of the coating, it was scratched using an AFM tip in contact mode under a constant force of approximately 2400 nN, until a uniform depth was achieved in the squared hole and no further material accumulation was observed on the surface. The height of this window was then taken as the film thickness. Nanoscope Analysis 1.5 software (Bruker) was used for image processing.

Roughness analysis was carried out from at least three different images (2.0 μm × 2.0 μm in size) and calculated as the root mean square (Rq) according to the Equation (1)

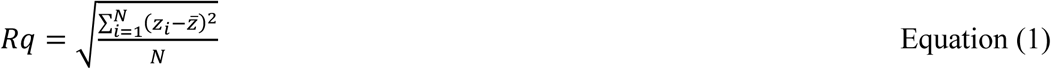

Where N is the number of points considered on the surface, z_i_ is the height of the point i on the surface, and z̅ is the average height of the N points.

### 2.8 Confocal microscopy

Confocal microscopy was used to reveal the amyloid-like structure of the aggregates. The PVC substrates covered by the BSA+DTT aggregates (BSA+DTT film) were immersed in a 2.06 × 10^−5^ M Tht aqueous solution for 15 min and then gently washed with Milli-Q water to eliminate the excess reagent. A PVC piece previously immersed in a BSA solution (without DTT) was used as a negative control The fluorescence intensity of the samples was then determined using a Carl Zeiss LSM 800 Microscopy. The measurement parameters were adjusted to 440 nm excitation wavelength and 485 nm emission wavelength, with a 20 μm pinhole. The images were processed using the Image J Software.

Also, confocal images were acquired using a live-dead fluorescence kit (Invitrogen). For this purpose, glass and PVC samples were first incubated for 24 h at 37°C with bacterial suspension at 10^5^ colony forming units (CFU)/mL and then washed to remove non adherent bacteria. Dye solutions were prepared by mixing 1µL of propidium iodine (PI) or Syto-9 in 3 mL sterile PBS. Finally, samples were stained for 15 min with Syto-9 solution, then washed and stained for 1 min with PI solution before imaging. Images were acquired using a PI channel with 561 nm of laser excitation at 0.4% and detection of emission at 618-700 nm Styt-9 chanel with 488 nm of excitation at 0.27% and 410-546 nm of emission detection, 52 μm pinhole. Both measurements were done with an EC plan - Neofluar 40x/0.75 M27 objective, and analyzed by ImageJ software.

### 2.9 Film stability assay

The stability of the film was evaluated by staining the film with Congo Red, which is specific to amyloid fibrils ^24^. A drop of 2.5 μM of Congo Red solution was poured on the BSA+DTT film on the PVC samples and left for 30 min. After that, the drop was removed and the samples were washed with Milli-Q water to eliminate the excess of reagent. The samples were immersed in ultrapure water and sonicated for 5 minutes to evaluate the film stability. The mechanic stability was tested by sticking and peeling off an adhesive tape on the film. After that, each sample was placed in a UV-vis Perkin-Elmer Lambda 35 spectrophotometer and the absorbance at 500 nm ^25^ was recorded before and after treatment (sonication or adhesive tape sticking).

### 2.10 Antibacterial assays

Microbiological assays were carried out using the protocol developed in our laboratory and previously reported for the evaluation of bacterial proliferation ^26^. Briefly, an overnight culture of *Staphylococcus aureus* (*S. aureus*, ATCC 25923) was prepared in nutrient broth (Biokar Diagnostics, France) at 37°C with agitation at 130 RPM. The bacterial suspension was diluted until ∼10^5^ - 10^6^ CFU.mL^−1^ and then the PVC substrates were vertically placed into the culture and incubated at 37°C for 2 h to allow bacterial adhesion. Once the incubation finished, the samples were gently washed with sterile water to eliminate weakly adhered bacteria to the surface. Subsequently, the samples were incubated in the sterile minimal medium GMP (glycine 133.2 mM, anhydrous dextrose 27.7 mM, and mannitol 27.5 mM, in phosphate buffer at pH 7.02) for 24 hours at 37 °C. After that, each sample was gently washed, sonicated in 1mL sterile phosphate buffer and then quantified by the plate count method. PVC and glass samples without the coating were employed as a negative control. To assess the effect of vancomycin on surface-adhered *S. aureus*, we first determined its minimum inhibitory concentration (MIC) against planktonic bacteria, which was found to be 0.625 µg/mL. Subsequently, varying sub-inhibitory concentrations of vancomycin (0.312 µg/mL, 0.156 µg/mL, and 0.078 µg/mL) were added to the GMP culture, and the same experimental procedures as previously described were applied. Among the tested concentrations, 0.078 µg/mL was selected as the working concentration, as it was the lowest dose capable of reducing the number of viable bacteria to about 10^5^ CFU/mm^2^ and CFU/mL for sessile and planktonic bacteria, respectively. This value is considered as a threshold for urinary tract infections and respiratory tract infections ^27–29^.

### 2.11 Cytocompatibility assays

HeLa (human cervical cancer) cells were cultured as sub-confluent monolayers in Eagle’s Minimum Essential Medium (EMEM) supplemented with 10% fetal bovine serum (FBS) and 1% penicillin/streptomycin at 37°C in a humidified atmosphere containing 5% CO₂. For cell seeding, the cells were detached using Accutase for 5 minutes, centrifuged, and the resulting cell pellet was resuspended in non-supplemented EMEM at the desired cell concentration. For the cell adhesion assay, the functionalized surfaces (previously washed and sterilized by 30 minutes of UV exposure) were placed in sterile 12-well plates containing 1 mL of medium and 150 µL of cell suspension (final cell concentration: approximately 2.5 × 10⁴ cells/mL). The fibronectin (FN) positive control was prepared by incubating the surfaces with 10 µg/mL of FN for 30 minutes at 37°C. The negative control was prepared by incubating PVC surfaces with 1% bovine serum albumin (BSA) for 30 minutes at 37°C. Both control surfaces were washed three times with sterile phosphate-buffered saline (PBS). The other functionalized surfaces were prepared using the same protocol as the experimental samples (e.g., BSA and BSA-DTT).

To quantify cell adhesion, the cells were washed once with sterile PBS, and the functionalized surfaces were removed from the 12-well plates and transferred to clean 24-well plates to ensure quantification of only the cells attached to the PVC surfaces. The 24-well plates were then placed in a −80°C freezer overnight. The next day, the plates were thawed, and each well was incubated with 500 µL of CyQUANT solution (prepared as a 20-fold dilution for cell lysis and 400-fold dilution of dye) for 5 minutes at room temperature (RT) on an orbital shaker. Subsequently, 200 µL of the CyQUANT solution was transferred to a 96-well plate, and fluorescence measurements were performed using a SPARK® Multimode Microplate Reader (TECAN) with excitation at 480 nm and emission measured at 520 nm.

For the toxicity assay, cells were seeded in 12-well plates without the functionalized surfaces and incubated overnight to establish a sub-confluent monolayer. The functionalized surfaces were then placed on top of the cell monolayers and incubated for 24 hours. After incubation, the surfaces were removed, the medium was aspirated, and the cells were washed with sterile PBS. After removing the washing liquid, the plates were frozen at −80°C overnight. Upon thawing, 700 µL of CyQUANT solution was added to each well and incubated for 5 minutes at RT on an orbital shaker. The solution was then diluted 1:10, and 200 µL was transferred to a 96-well plate. Fluorescence intensity was measured as described previously, using excitation at 480 nm and emission at 520 nm.

### 2.12 Statistical analysis

All experiments were performed independently, mostly in triplicate. When replicates were not available, data are presented as individual values. Microbiological and cytocompatibility assays were conducted in three independent experiments. Results are expressed as mean ± standard deviation. Significant differences were determined using one-way analysis of variance (ANOVA) followed by Tukey’s post hoc test (p < 0.05).

## 3 Results

### 3.1 Albumin-based amyloid-like aggregates formation at physiological conditions

Amorphous aggregates like amyloids were prepared by reducing disulfide bonds in BSA by dithiothreitol (DTT), which destabilizes the native protein structure and thereby promotes aggregation. The reaction converts alpha-helical structures of the native protein to beta-sheets which are present in the new structure obtained. Upon incubation, these beta-sheets stack on each other to form aggregates like amyloids ^30^. Over time, the macroscopic characteristics of the BSA+DTT dispersions change, becoming increasingly turbid, as previously reported for BSA amyloid-like solutions ^20^. In order to confirm the amyloid-like structure of the aggregates, fluorescence spectrophotometry assays were carried out by adding ThT to the aggregates solution. ThT has been used as a gold standard probe to detect amyloid fibrils as it binds to the channels running parallel to the long axis of the fibrils, giving rise to its characteristic emission fluorescence peak around 485 nm upon excitation at 450 nm ^31^. During the adsorption of thioflavin on the surface of the aggregates, an internal charge transfer occurs due to ThT adopting a planar conformation when interacting with amyloid structures. This change in the conformation of ThT from a non-radiative state to a locally excited state results in the fluorescence emission^32^. Figure 1a shows the fluorescence intensity profile after the addition of ThT to BSA+DTT. For comparison, the spectra of BSA+DTT (without ThT) and BSA+ThT (without DTT) were also recorded. In the presence of DTT, the emission of ThT increases drastically with time without lag phase in the aggregation process, which is consistent with previous reports ^20^. The increment in the intensity of the ThT emission with time reflects the increase in the aggregates concentration as these variables are directly proportional. The native protein in the absence of DTT does not show any change in the ThT emission, indicating that under these conditions the protein has not undergone any amyloid nucleation. Transmission electron microscopy (TEM) images (Figure 1b) reveal that the BSA+DTT aggregates include some fibrillary structures, consistent with typical amyloid formation. These fibrils appear intertwined, forming regions of higher aggregation (darker areas in the image). From this entanglement, the fibrillar clusters tend to develop branched structures that spread across the resin surface of the TEM grid. Since most observations reveal amorphous aggregates composed of intertwined fibrils, we refer to them broadly as aggregates rather than amyloid fibrils throughout this study. It is worth noting that the temperature used for depositing the BSA+DTT aggregates onto the TEM grid (4 °C) may slightly influence their morphology.

**Figure 1.**
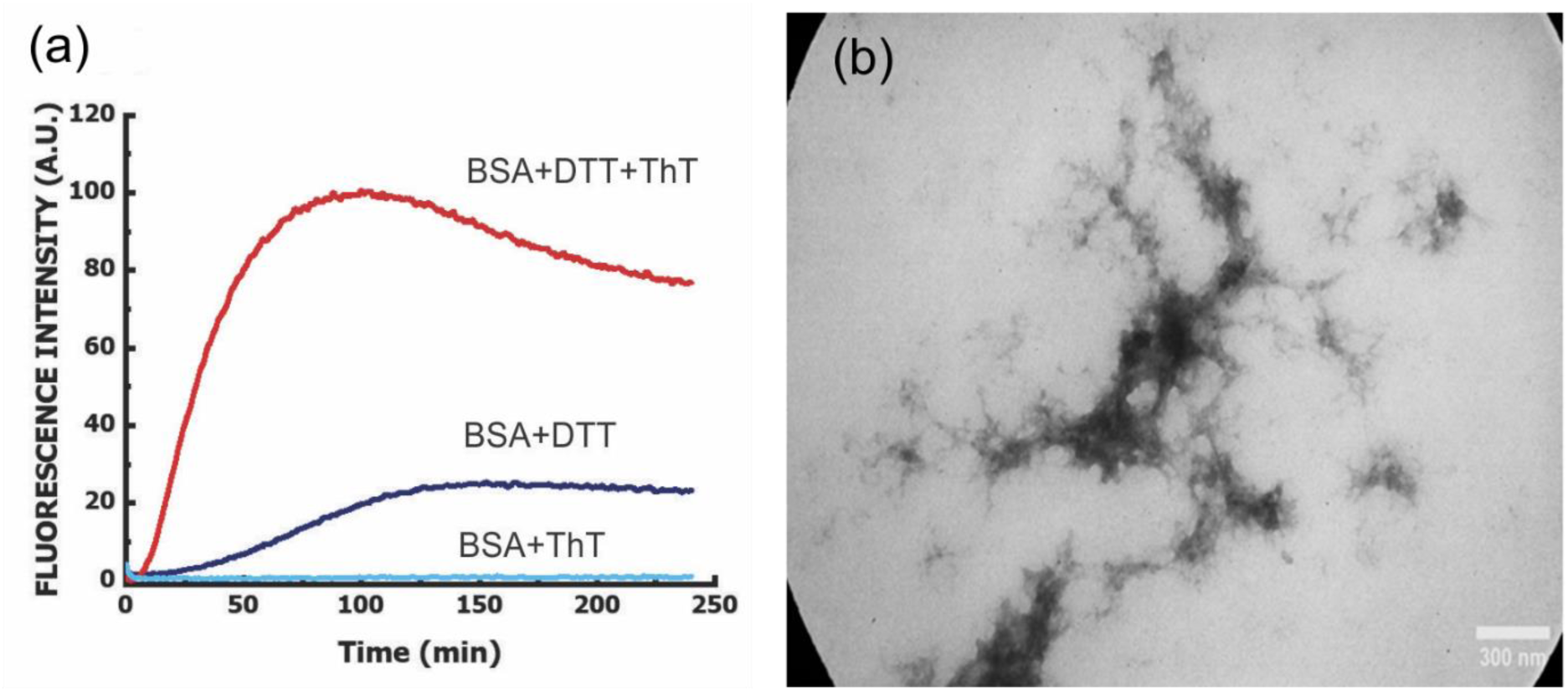
(a) Fluorescence spectroscopy of BSA+DTT before and after ThT addition. The spectrum corresponding to BSA treated with ThT is included for comparison. (b) TEM image of representative BSA+DTT aggregates.

### 3.2 Formation of an amyloid-like albumin coating on glass and PVC surfaces

First, we explored the coating of glass as a model surface to study the formation of amyloid aggregates hydrogel films. Then we extended the study to PVC surfaces as a biomaterial of interest, since various medical devices susceptible to bacterial colonization (catheters, medical tubes such as dialysis and endotracheal tubes, etc.) are made of this material ^22^. We used the immersion of the samples into the BSA+DTT solution as a simple method to coat the surfaces. The modified surfaces were then characterized with a multi-technique approach. As a first approximation, the surface staining with Congo Red allowed to confirm the formation of a coating on the surface. Congo Red is commonly used to identify amyloid fibrils in biological samples ^24^, as the dye binds to beta-sheet structures through several interactions, including van der Waals, hydrogen bonding, and hydrophobic-hydrophobic interactions. The binding of Congo Red to amyloid fibrils is highly selective; therefore, staining the samples exposed to the BSA+DTT solution would indicate the formation of an amyloid-like film on the surface. Figure 2 shows Congo Red-stained samples as a function of the immersion time of glass and PVC samples in BSA+DTT solution. As control, samples exposed to just BSA solution were tested. It can be noticed that the red colour intensity in BSA+DTT exposed samples visibly increases as the immersion time does, indicating not only that the amyloid-like film has been formed, but also that the film can cover the surface with a rise in the film thickness with time. Conversely, samples exposed to BSA remain uncoloured, meaning no aggregates were formed on the surface. Since 6 h immersion leads to the thickest film, we have chosen this exposition time for the next experiments.

**Figure 2.**
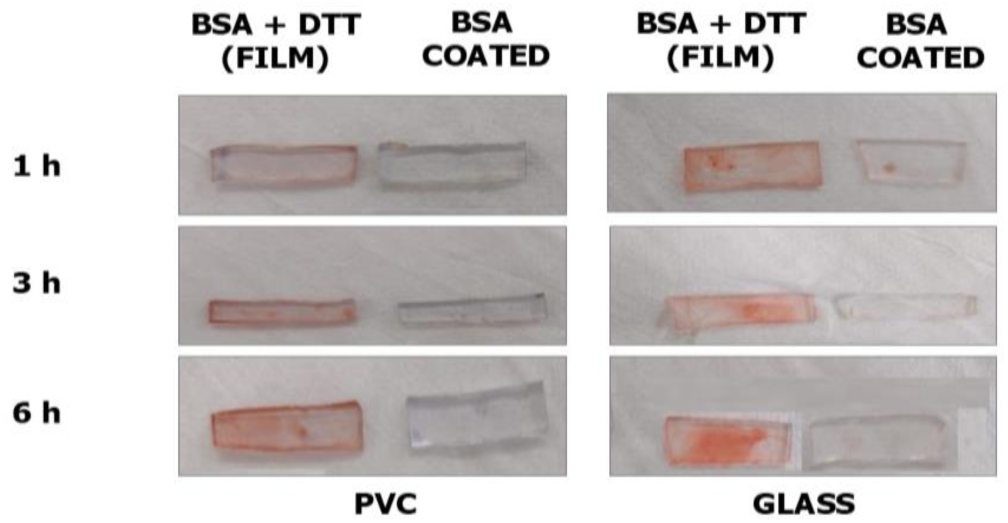
Congo Red staining of the amyloid-like film on glass and PVC surfaces as a function of the exposition time to BSA+DTT dispersion. As control, samples exposed to BSA were also assayed.

FTIR is a commonly used technique for characterizing the structure and composition of amyloid fibrils deposited on surfaces. FTIR analysis of amyloid fibrils provides information on the secondary structure, conformational changes, and molecular interactions that occur during amyloid formation. We have used ATR-FTIR to characterise the albumin amyloid films in both, in solution and deposited on PVC (Figure 3). The spectrum of the solution allows us to identify the expected bands. The characteristic band of BSA deposited on PVC was identified and a shift towards a lower wavelength for the PVC surfaces covered with the BSA+DTT film was found. The FTIR spectra of BSA in its soluble form typically show distinct bands at approximately 1650 cm^-1^ and 1550 cm^-1^, which correspond to amide I and amide II vibrations, respectively ^33^. These bands are characteristic of the protein’s secondary structure, which is predominantly composed of random coil conformations. In contrast, amyloid-like aggregates formed from BSA show different FTIR spectra. The FTIR spectra of amyloid fibrils typically show a sharp band at 1620 cm^-1^, characteristic of beta-sheet conformations. The presence of this band (Figure 3) indicated the formation of beta-sheet amyloid fibrils, which is a hallmark of amyloid structures ^33^. Additionally, the FTIR spectra of amyloid fibrils may also show a shift in the position of the amide I band from approximately 1650 cm^-1^ in native BSA to around 1640 cm^-1^ in the amyloid form. This shift is indicative of a protein change and is often used as an indicator of amyloid formation.

**Figure 3.**
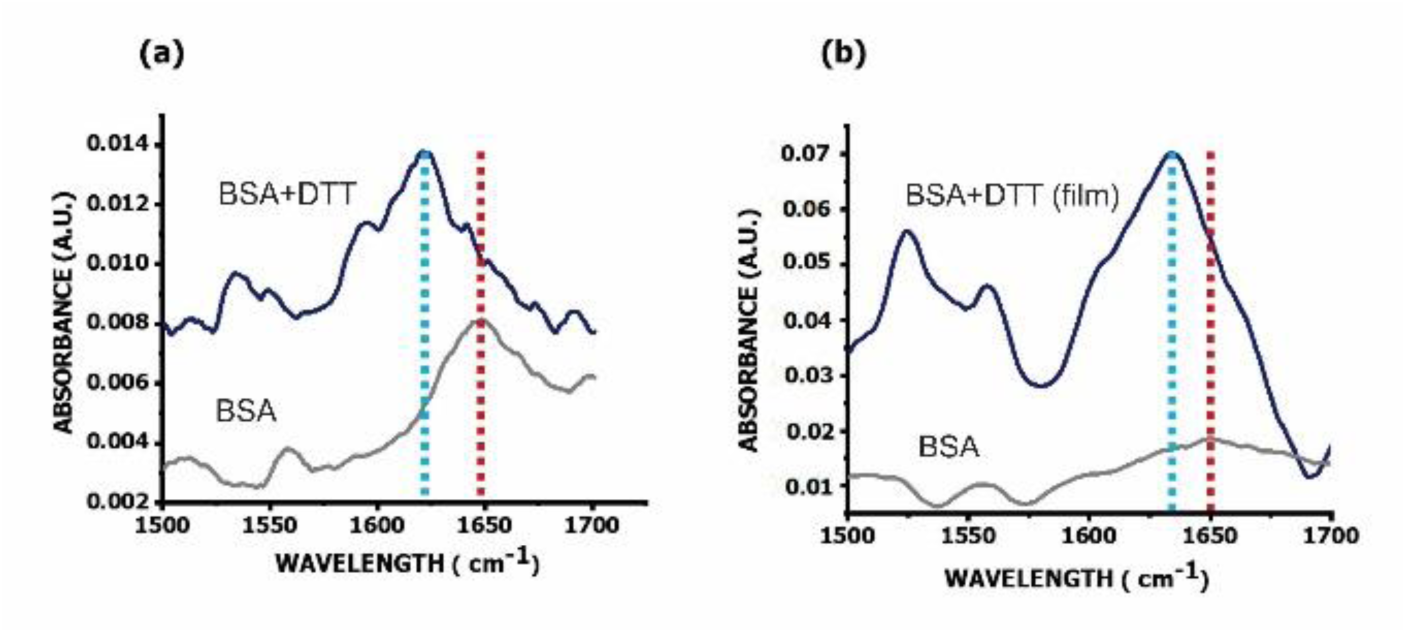
ATR-FTIR of BSA and BSA+DTT (a) in solution; (b) film on PVC, showing the typical shift in the amide I band corresponding to BSA (red dotted line, 1650 cm^-1^) when amyloid structures are formed (blue dotted line, 1630 cm^-1^).

The wetting properties of the film were evaluated using contact angle measurements (Figure 4). Glass is a highly hydrophilic surface (θ = 14.7 ± 0.4), whereas PVC is significantly less hydrophilic (θ = 80.0 ± 0.6). The presence of the film alters the contact angle, resulting in values of 43 ± 7 for glass and 65 ± 2 for PVC. These variations in contact angle for the BSA+DTT film, depending on the substrate, likely reflect the macroscopic nature of the measurement. This suggests that the film does not entirely cover the surface, allowing the underlying substrate properties to influence the results.

**Figure 4.**
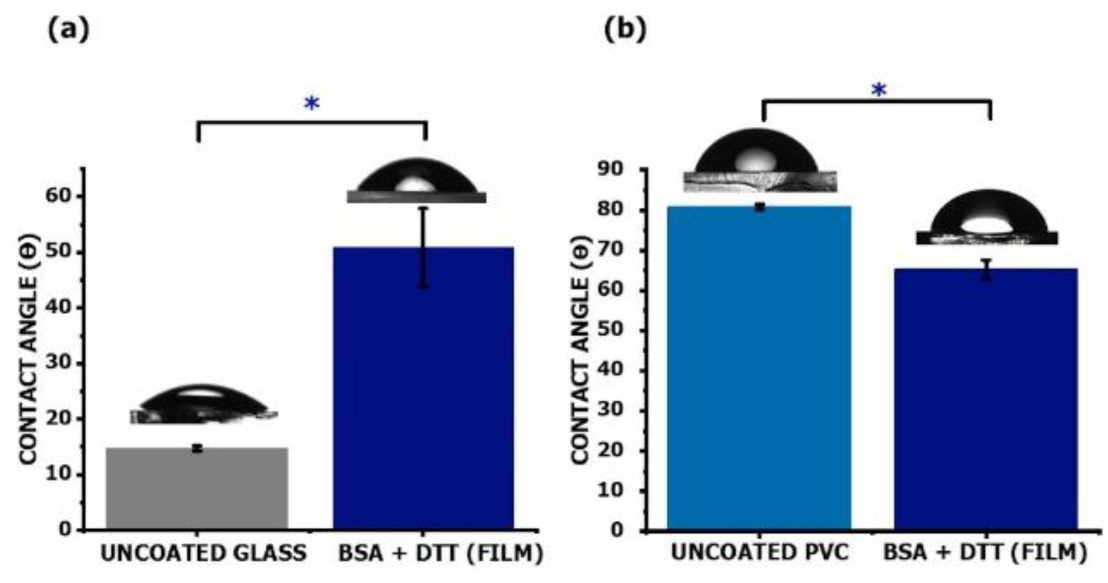
Contact angle for the uncoated substrate and the BSA+DTT amyloid-like film. (a) Glass; (b) PVC

ThT-stained samples were also analyzed by confocal microscopy, measuring the fluorescence intensity on different regions of each sample (Figure 5). The BSA+DTT-modified surfaces exhibited higher fluorescence intensity compared to those modified with BSA alone. The average fluorescence intensity measured along the lines shown in Figures 5A(b) and 5B(b) was 1.0 ± 0.1 for BSA-coated glass and 1.0 ± 0.5 for BSA-coated PVC. In contrast, the fluorescence intensity for the BSA+DTT film was 2.35 ± 0.05 on glass and 8.2 ± 0.1 on PVC. Notice that Figure 5 A(b) suggests that the BSA+DTT film does not uniformly cover the glass surface. This observation aligns with the lower contact angle measured for the film on glass compared to PVC. Since the contact angle represents an average measurement over the surface, the lower value on glass may reflect the influence of uncovered regions. On the other hand, the lower fluorescence intensity recorded for the film on glass could be related to the lower thickness of BSA+DTT film on this substrate (see Figure 7A). In this regard, the nature of surfaces modulates de kinetics of amyloid formation and aggregation ^34^. Various protein-substrate interactions, influenced by the predominant functional groups exposed by the aggregates (aromatic rings, hydroxyl groups, carboxyl groups, amines, etc.), may play a key role in driving adsorption. Multiple binding sites, often involving hydrogen bonding, hydrophobic forces, and electrostatic attractions, among others, can be formed between the film and the underlying substrate ^35^, contributing to the stable interfacial adhesion of the thin film across the surfaces. More hydrophilic surfaces lead to a lower BSA adsorption ^36^. Thus, a prolonged residence time might be necessary to improve the alignment of contact points between albumin and hydrophilic substrates, ensuring optimal interaction even under favourable electrostatic conditions ^37^. Therefore, considering that glass is more hydrophilic than PVC, it is likely that applying the same immersion time to both substrates results in a lower adsorption of BSA+DTT aggregates on glass.

**Figure 5.**
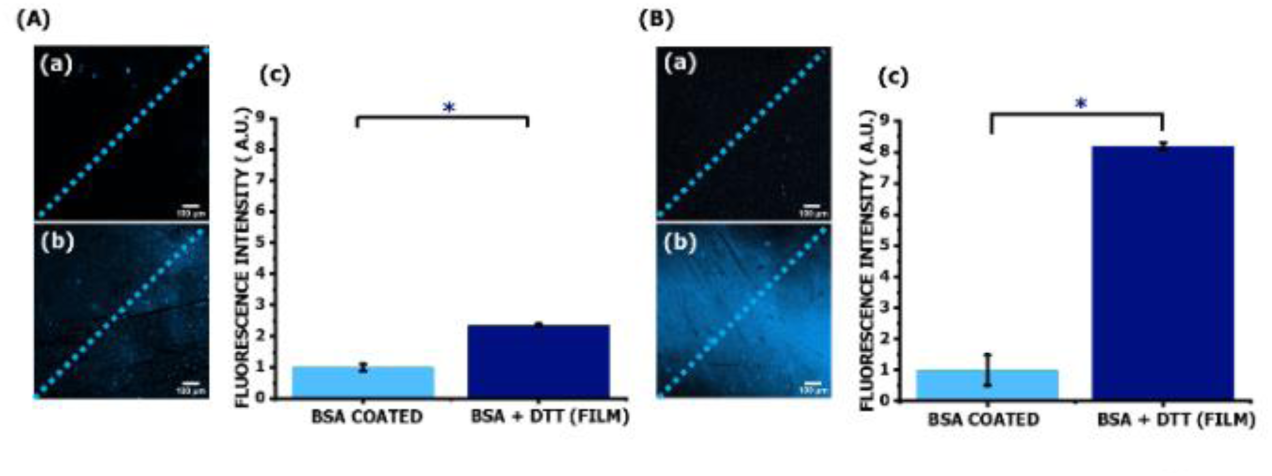
Confocal microscopy imaging (A) Glass samples. (B) PVC samples. For both substrates: (a) ThT-stained BSA coating (b) ThT stained BSA + DTT (film) coating. (c) Fluorescence intensity mean value for the different substrates along the lines in the images shown in (a).

To further characterize the amyloid-like BSA coating, AFM imaging of the surfaces was performed (Figure 6). The film on both, glass (Figure 6A) and PVC (Figure 6B), consists of a uniform layer of amyloid aggregates, on which higher, irregular branched structures can be found in some regions. This description is supported by the surface roughness Rq (Table 1) measured on the bare substrates and the underlying film (avoiding higher branched structures), with values that remain consistent across different regions of the same substrate. As expected, uncoated glass substrates are smoother (Rq = 0.34 ± 0.04 nm) than uncoated PVC (Rq = 4.4 ± 0.9 nm). However, after coating, the roughness increases by 400% for glass and 36% on PVC. Notice that although PVC is 13 times rougher than glass, the Rq value for the BSA+DTT film on PVC is only 3.5 times higher than that on glass. This discrepancy suggests a non-conformal deposition, where the material accumulates unevenly, with greater build-up in some areas and less in others. The lack of proportionality between the Rq values and the roughness of the bare substrates further supports this observation. The film thickness on both substrates was estimated by scratching it with an AFM tip (Figure 7). The height results in 202 ± 32 nm for glass and 1.3 ± 0.7 µm for PVC.

**Figure 6.**
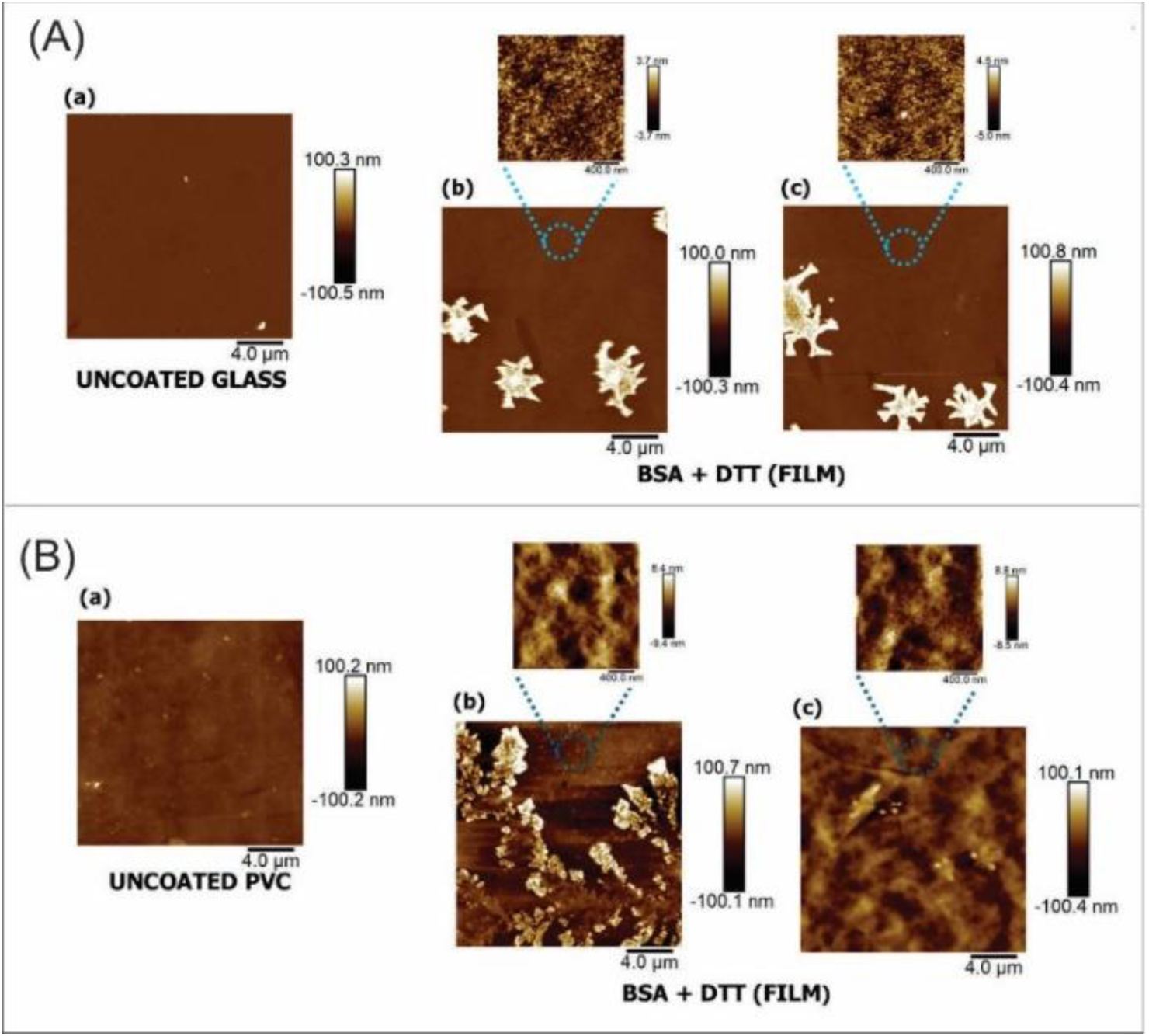
AFM images (20 µm x 20 µm) of (A) glass and (B) PVC. For both samples: (a) uncoated surface; (b) and (c) different regions of the BSA+DTT film; the zooms show details of the underlying layer.

**Figure 7.**
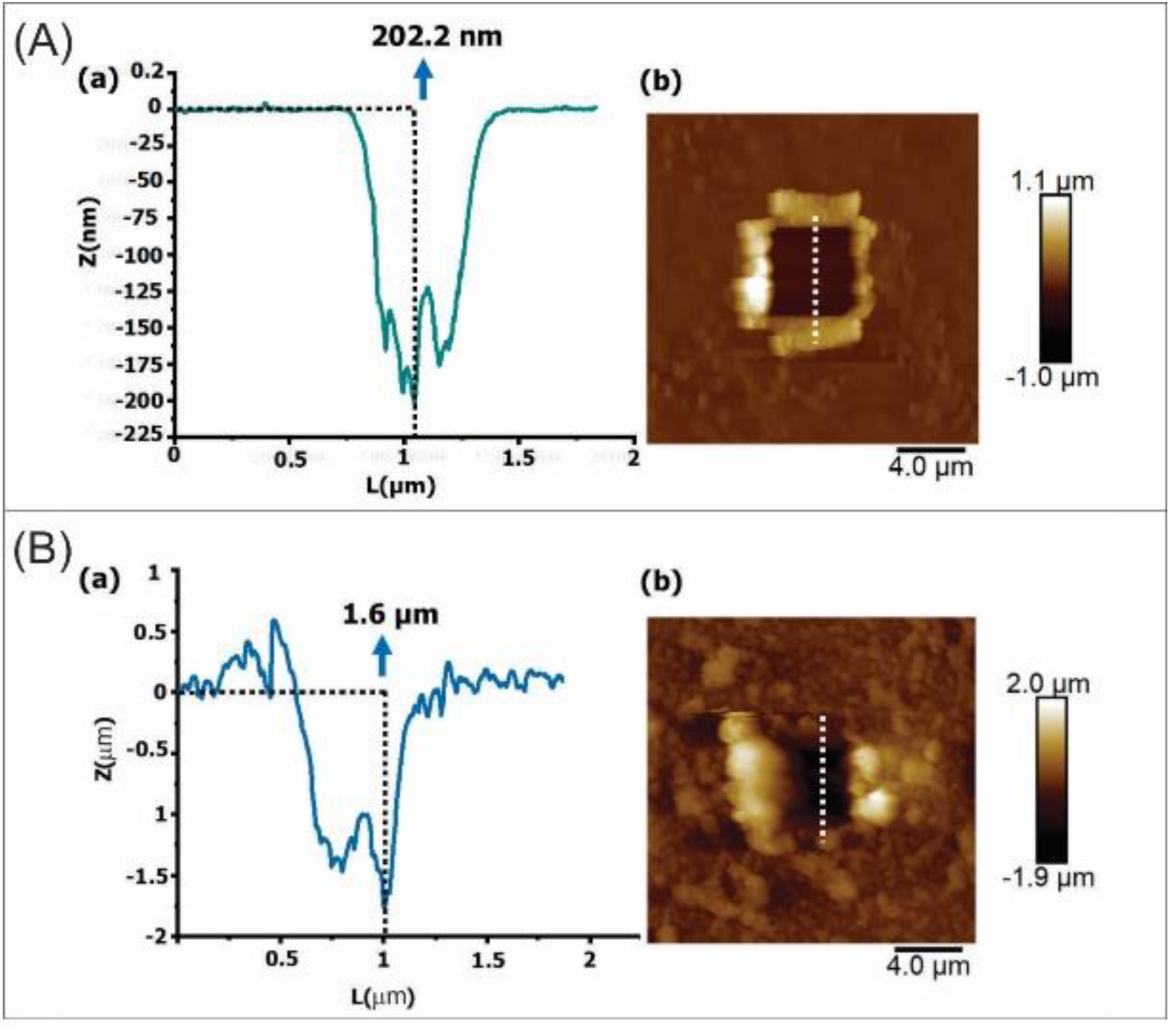
Film thickness measured by scratching the film with an AFM tip. (A) glass, (B), PVC. For both samples: (a) cross section along the line shown in the AFM image (b). A representative individual image and value for the height of the BSA+DTT film on each substrate is shown.

**Table 1.** Root mean square (Rq) measured on 2.0 μm × 2.0 μm AFM images.

|  | Rq (nm) |
| --- | --- |
| Uncoated glass | $0.34 \pm 0.04$ |
| BSA+DTT (film) on glass | $1.7 \pm 0.1$ |
| Uncoated PVC | $4.4 \pm 0.9$ |
| BSA+DTT (film) on PVC | $6 \pm 1$ |

### 3.3 Biocompatibility of BSA+DTT films

The application of this antimicrobial coating on a biomedical device requires the analysis of the corresponding biocompatibility and cytotoxicity. ISO standards stipulate the requirements for the biological evaluation of medical devices. ISO 10993 (Biological Evaluation of Medical Devices) prescribes HeLa cells as one of the adequate cells for general biocompatibility of any biomaterial. The method to quantitatively evaluate biocompatibility and cytotoxicity of the coating was performed using CyQuant®, Cell Proliferation Assay. CyQuant® assay is a rapid method for monitoring cell adhesion and has the sensitivity to measure small numbers of adherent cells. CyQuant®GR dye is used to selectively label cellular DNA post experimentation and provides a rapid readout of adherent cell number without the time-consuming pre-labelling procedure.

The biological evaluation was explored by means of two different types of assays: a cell adhesion and a toxicity assay. For the cell adhesion experiment, the cells were seeded on top of the different functionalized surfaces. A negative (BSA 1%) and a positive control (FN 10 μg/mL) were also tested. BSA 1% is commonly used as a negative adhesion control. BSA physically prevents other proteins (like fibronectin or vitronectin) from binding. These extracellular matrix proteins are essential for integrin-mediated cell adhesion, and without them cells will not attach to BSA-coated wells. As expected, after 1 hour the cells could successfully spread on the surface with fibronectin coating (positive control). On the surface coated with BSA 1%, which is commonly used to hinder cell adhesion on surfaces, the cells were not able to adhere. In order to check the adhesion of cells on the functionalized surfaces, we have used not only the surface with the amyloid-like BSA coating but also the surfaces coated only with BSA at the same concentration used to prepare the coating. All the tested samples hindered the adhesion of cells on the surfaces (Figure 8A).

**Figure 8.**
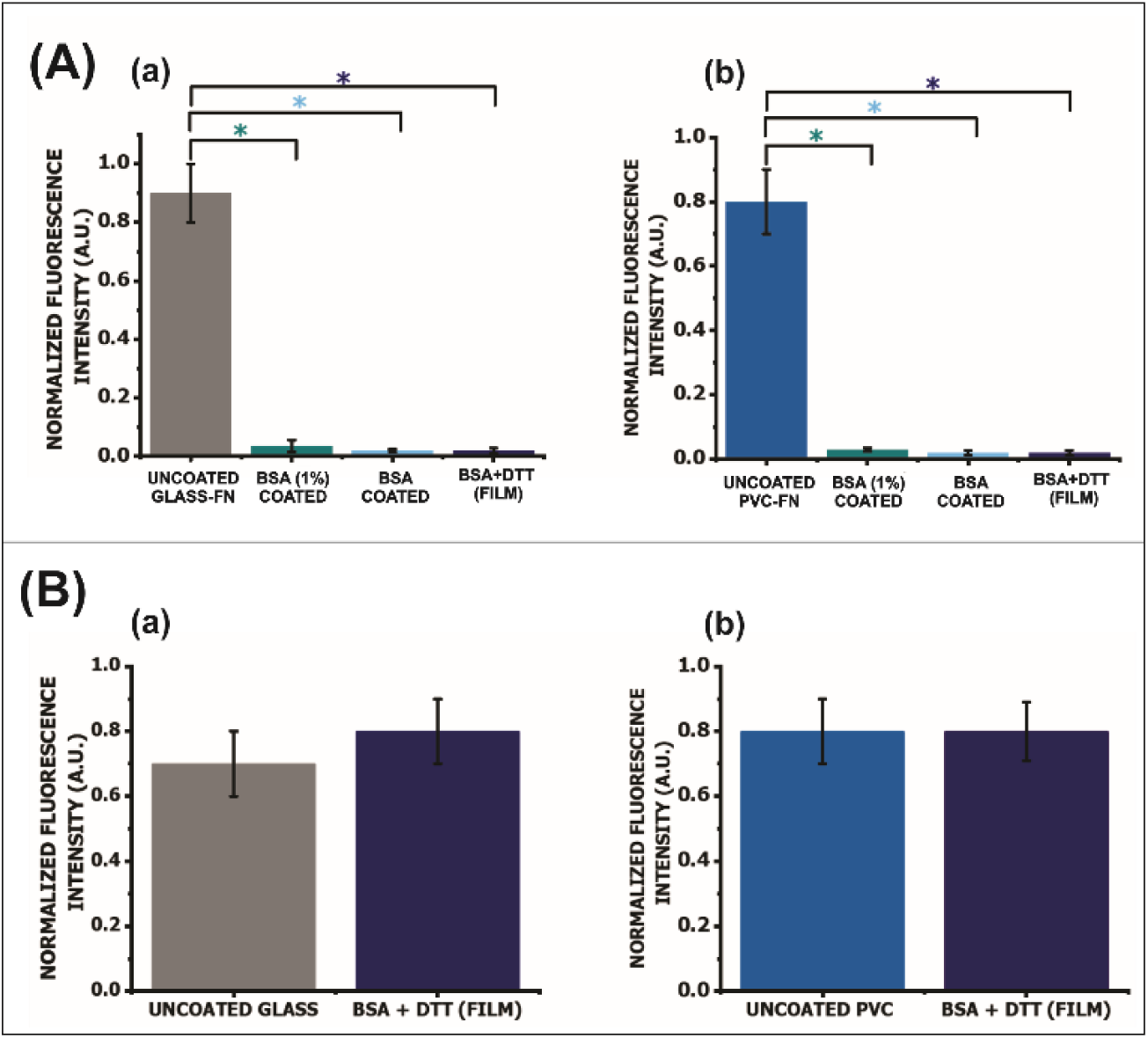
(A) HeLa cells adhesion on the amyloid aggregates film. (B) Cytotoxicity of the amyloid aggregates film on HeLa cells. For both figures, (a) on glass; (b) on PVC. (*) indicate significant differences.

We will discuss these results in relation to the physicochemical parameters measured for the BSA+DTT films coated on glass and PVC surfaces. Cell adhesion depends on various surface characteristics, including wettability, roughness, and surface chemistry. The interplay among these parameters determines the extent to which cells adhere to a given surface. Most mammalian cells typically attach and proliferate on surfaces that are moderately hydrophilic, rather than on those that are extremely hydrophilic or highly hydrophobic. While hydrophilic surfaces can promote cell differentiation, superhydrophobic surfaces may disrupt fibronectin conformation, thereby impairing cell adhesion ^38^. It has been suggested that surfaces with contact angles between 40° and 70° are optimal for promoting cell adhesion ^38^. However, despite the BSA+DTT films exhibiting contact angles within this favourable range, they behave as anti-adhesive coatings. This observation indicates that wettability is not the primary factor governing the anti-adhesive behaviour of the films.

Surface topography also plays a crucial role in modulating cell behavior across different length scales. Macroscale features (>100 μm) influence cell colonies or populations; microscale structures (0.1–100 μm) can affect individual cells by guiding their shape, orientation, and migration; and nanoscale textures (1–100 nm) can interact with membrane receptors, thereby modulating signaling pathways and cellular responses ^39^. However, discrepancies in the literature regarding the topographical features that promote cell adhesion suggest that the underlying mechanisms remain poorly understood ^40^. Previous studies have shown that moderate nanoscale roughness can enhance cell adhesion by providing physical cues that promote the formation of focal adhesions. For example, Hou *et al.* ^40^ designed surfaces with a roughness gradient ranging from nanoscale to microscale and found a linear correlation between surface roughness and cell spreading. Interestingly, although the cell spreading area increased linearly with roughness, focal adhesion formation, filopodia extension, and cellular tension followed a biphasic trend, peaking at an intermediate roughness of approximately 200 nm.

In our study, the surface roughness of all samples is relatively low (less than 8 nm, see Table 1). The film roughness on glass and PVC substrates is comparable, differing only by a factor of 3.4. Therefore, it is unlikely that roughness is the primary factor responsible for the inhibition of cell adhesion. We therefore propose that the anti-adhesive properties of BSA+DTT films are primarily determined by the chemical nature of the coating. As previously noted, BSA is a well-established passivating agent known to inhibit cell adhesion. It achieves this by physically preventing the adsorption of extracellular matrix proteins such as fibronectin and vitronectin, which are crucial for integrin-mediated attachment. In the absence of these proteins, cells are unable to effectively adhere to the surface ^41,42^. However, BSA amyloid-like aggregates may exhibit a distinct behavior. Diaz *et al*. ^20^ thoroughly characterized physically cross-linked amyloid-like BSA fibrils and demonstrated that these structures can inhibit cell adhesion by forming a dense, cross-linked fibrillar network. Rich in β-sheet content, this network lacks the molecular cues necessary for cell binding, thereby preventing both cell attachment and the adsorption of adhesion-facilitating proteins. Accordingly, the amyloid-like BSA aggregates maintain their anti-adhesive properties, effectively inhibiting cell attachment. To confirm that the coating is non-toxic, we have designed a different experiment. A monolayer of HeLa cells was grown on a well plate as control (PS) and the surfaces functionalized with the different coatings were placed on top of the monolayer, directly in contact with cells growing on the culture medium. The cells were then incubated with the functionalized surface on top. During incubation, if leachable toxic molecules from the coating are released, these compounds could diffuse into the culture medium and contact the cell layer. Reactivity of the test sample is indicated by malformation, degeneration and lysis of cells around the test material. The obtained qualitative and quantitative results demonstrate that the hybrid coating has no toxic element that could disrupt the cell layer (Figure 8B).

For a material to be clinically applicable, it must exhibit excellent biocompatibility, meaning it should perform its intended function without causing harmful local or systemic effects. The release of toxic substances at the implant site can lead to cell death and subsequent tissue damage. However, regardless of the deposition conditions, the films used in this study showed no toxic effects on HeLa cells. From the cell adhesion and cytotoxicity studies, we could confirm that the amyloid-like BSA coating is biocompatible with anti-adhesive characteristics.

### 3.3 Antibacterial properties of BSA+DTT films

The biocompatibility of the BSA + DTT coating has been confirmed; the next step is to assess its antibacterial properties. The question that arises is: Why use amyloid-like aggregates to inhibit cell fouling instead of just BSA (Figure 8)? Our results demonstrate that surfaces coated with BSA alone do not inhibit bacterial adhesion and may even promote it, potentially increasing the risk of infections (Figure 9).

**Figure 9.**
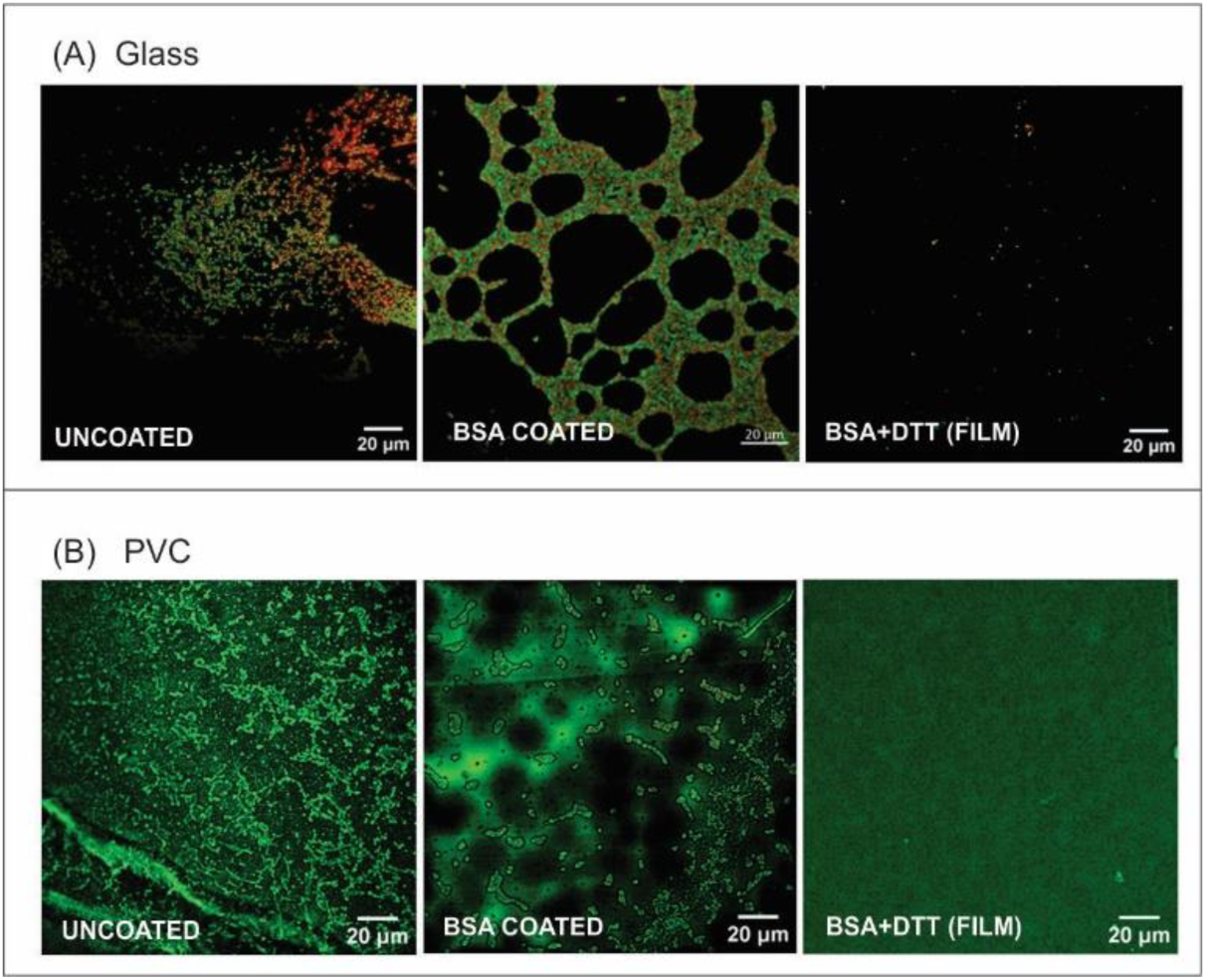
Confocal microscopy images of bacterial attachment after 2 h exposition to *S aureus* culture. (A) Glass; (B) PVC. It should be noted that, under the measurement conditions, the PVC substrate exhibited a green autofluorescence, making it difficult to distinguish bacteria adhered to the surfaces. Therefore, the background of the images has been removed to improve visualization.

The effectiveness of the BSA+DTT-modified surfaces in inhibiting *S. aureus* growth and proliferation was evaluated. *S. aureus* is a major pathogen in catheter-related infections, notably responsible for 34.2% of bloodstream infections reported in U.S. dialysis facilities in 2020 ^43^. It is a leading cause of central line-associated bloodstream infections and is frequently linked to high morbidity and mortality, with rates reaching up to 40% in cases of *S. aureus* bacteremia ^44^. Also, *S. aureus* is a significant pathogen in ventilator-associated pneumonia (VAP) and infections related to endotracheal tubes (ETTs) ^45^. In studies analysing endotracheal aspirates, *S. aureus* was identified as the most common pathogen, present in 56.8% of samples, and was associated with a higher likelihood of developing VAP ^46^. Furthermore, biofilm formation by *S. aureus* on ETTs contributes to its persistence and resistance to treatment, complicating infection management. These findings highlight its critical role in tubing-associated infections and the importance of effective prevention strategies.

As an initial step, we investigated whether the anti-adhesive properties of the film observed with HeLa cells also extended to bacterial adhesion.

Confocal microscopy was employed to assess the attachment of *S. aureus* on the modified surfaces following exposure to the bacterial culture. Bare substrates were used as controls. As shown in Figure 9, the BSA+DTT film effectively inhibited bacterial attachment on both types of substrates, as only a few bacteria can be seen on these surfaces. Notably, surfaces modified with BSA alone failed to prevent *S. aureus* adhesion, underscoring the critical role of amyloid-like aggregates in achieving anti-adhesive properties.

To evaluate bacterial behaviour on the modified surfaces, an experimental model was employed based on the quantification of viable surface-associated bacteria using the serial dilution method. To this end, after bacterial attachment, the growth and proliferation on the PVC and PVC-modified surfaces were assessed in sterile GMP medium over a 24-hour period, both in the presence and absence of vancomycin. The results presented in Figure 10 confirm the anti-biofouling properties of the BSA+DTT films, as the number of viable sessile *S. aureus* cells decreased three-log compared to bare PVC, used as the control (Figure 10a), resulting in a bactericidal surface. Similarly, the number of sessile bacteria that detached from the surface—likely during the early stages of exposure to the sterile medium—and transitioned to a planktonic phenotype was reduced four-log relative to the control (Figure 10b). Notably, the addition of vancomycin to the sterile medium resulted in the complete eradication of both sessile and planktonic *S. aureus* on the BSA+DTT-coated substrates. This indicates a synergistic effect ^47^ between the BSA+DTT (film) and vancomycin, as vancomycin alone reduced the number of viable bacteria on uncoated PVC by only two-log for sessile and one-log for planktonic forms. Importantly, this effect was achieved using a vancomycin concentration that was eight times lower than the MIC.

**Figure 10.**
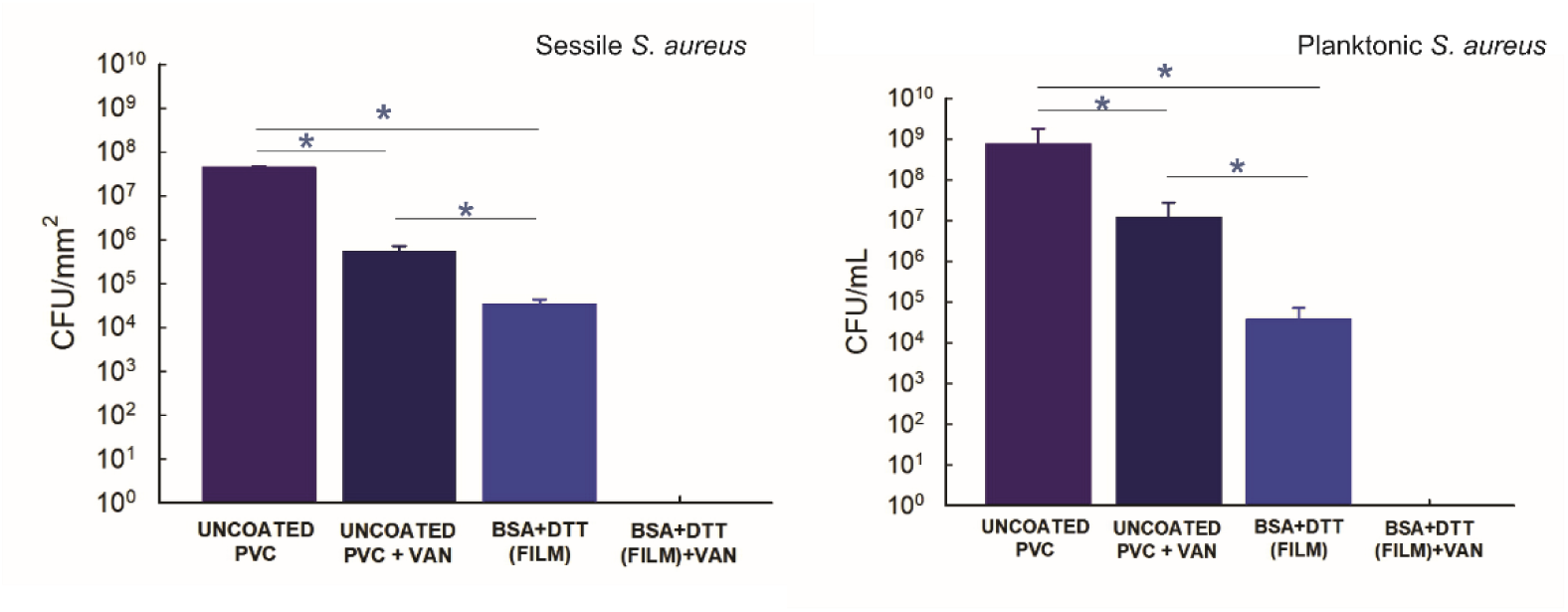
Viable bacteria after 24 hours of growth and proliferation in sterile GMP medium with and without vancomycin (VAN). (a) Sessile bacteria attached to each surface; (b) Planktonic bacteria in each supernatant. Asterisks (*) indicate statistically significant differences. Note that CFU counts corresponding to BSA+DTT (FILM)+VAN were not included in the statistical analysis, as the zero value observed for these populations indicates that CFUs were below the limit of detection.

Medical tubing is widely employed to treat various pathologies and sustain life. Common applications include feeding and infusion tubes, tubing for hemodialysis and peritoneal dialysis, urinary catheterization, and assisted ventilation via endotracheal tubes, among others. However, tubing systems can fail due to multiple factors during clinical procedures. Complications such as thrombosis, bacterial contamination, blockages, or improper insertion can compromise device function and negatively impact patient health. Failure of drainage catheters, drug delivery devices, and other tubing systems often results from fibrous capsule formation, scar tissue blockage, or fibrous encasement ^48–51^. Notably, the formation of bacterial biofilms not only promotes infection but also contributes to tube blockage through the crystallization of biofilms ^52,53^. Importantly, we demonstrate the synergistic effect of the coating upon vancomycin treatment, which is clinically relevant as it leads to complete bacterial eradication. Although the amyloid film alone exhibits anti-biofouling properties and reduces bacterial adhesion compared to the control, the remaining bacterial load on the surface still reaches levels associated with infection risk in indwelling medical devices. ^29^. Remarkably, this synergistic effect is achieved using a vancomycin concentration eight times lower than the MIC, highlighting the potential of the coating to enhance antibiotic efficacy while minimizing drug usage. Therefore, the results presented in this study may offer a potential strategy to mitigate tube malfunction by reducing both cell and bacterial adhesion, while simultaneously enhancing the efficacy of conventional antibiotic treatments.

### 3.4 Stability of the BSA+DTT films

Finally, since the primary goal of the BSA+DTT films is to provide coverage for infection-prone PVC medical devices, mechanical stability is crucial to prevent detachment during insertion. Additionally, stability in a liquid environment is important when the film is subjected to stress. Therefore, we have investigated the BSA+DTT film stability in aqueous media (sonication in ultrapure water) as well as their mechanical durability (peeling off with adhesive tape) (see section 2.12). To this aim, the samples were dyed with of Congo Red and the absorbance was measured before and after the stability challenge. Figure 11 shows that, although the absorbance due to Congo Red staining slightly decreases, the film remains on the surface for both, sonication in ultrapure water and sticking and peeling off an adhesive tape. These results are consistent with the relatively high force needed to sweep the film with an AFM tip (about 2400 nN, see Section 2.9) which indicates the strong adhesion to the surface. On this regard, it should be noted that deciphering the precise molecular mechanisms of protein-surface interactions remains a significant challenge. Establishing universal principles for predicting them is highly complex due to the multitude of physical and chemical factors involved, which are often system-specific.

**Figure 11.**
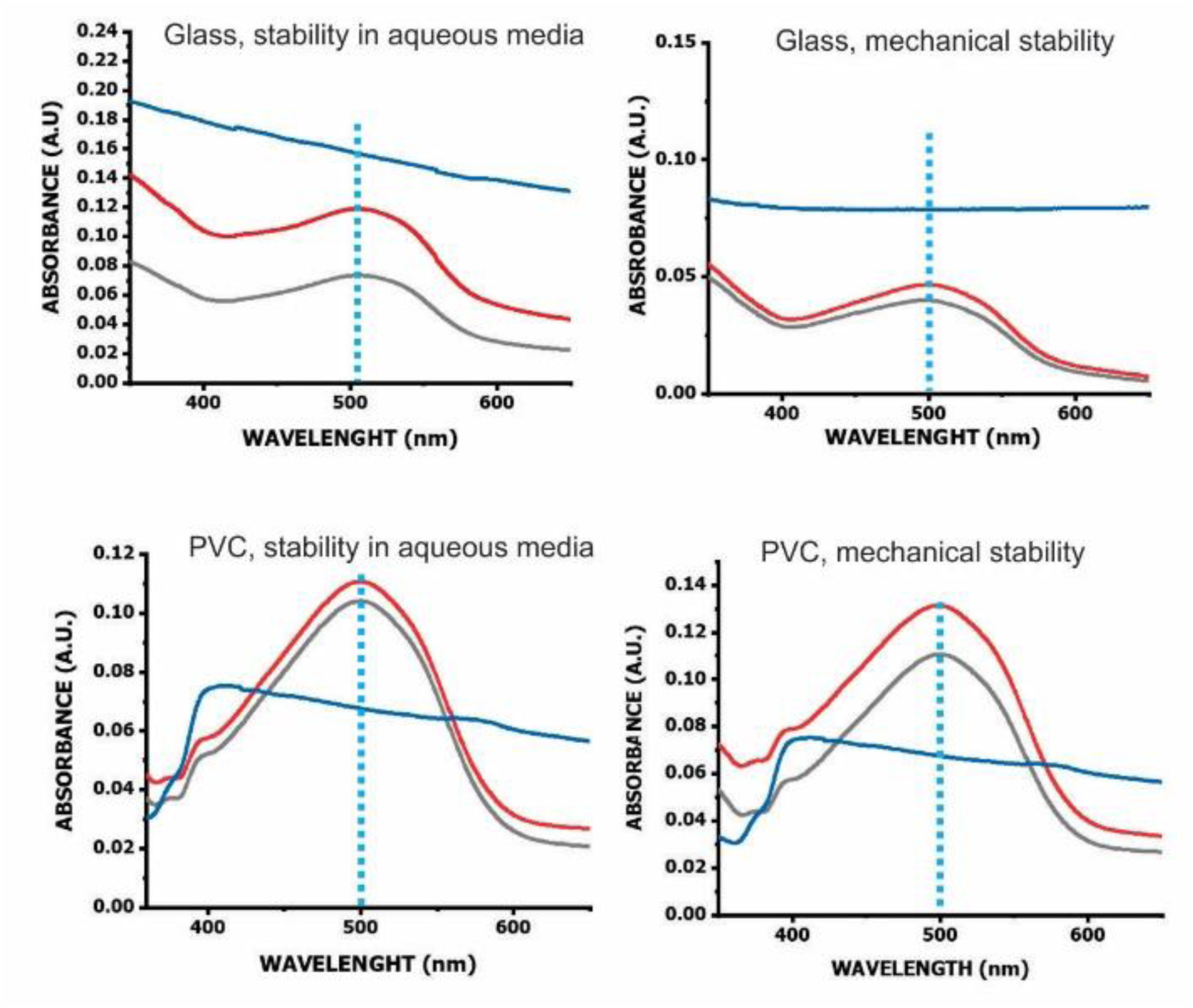
UV-vis spectra of Congo Red-stained BSA+DTT film on glass and PVC before (red line) and after (grey line) sonication in ultrapure water (stability in aqueous media) and before (red line) and after (grey line) sticking and peeling off an adhesive tape (mechanical stability). For comparison, the spectrum of BSA+DTT film without dying (blue line) is also included.

## 4 Conclusions

This study explores an alternative synthesis route for generating functional anti-biofouling coatings from bovine serum albumin (BSA) amyloid-like aggregates for medical tubing applications. While previous studies have reported the anti-biofouling properties of BSA amyloid coatings, here we demonstrate that this alternative preparation method preserve the main characteristic physicochemical features and anti-adhesive performance of the previously described coatings. The BSA amyloid-like coatings exhibited significant resistance to bacterial adhesion, particularly against *Staphylococcus aureus*, a major pathogen in device-related infections. Importantly, our findings reveal a synergistic effect between the amyloid-like coating and vancomycin treatment, resulting in complete bacterial eradication on the coated surfaces. Notably, this effect was achieved using a vancomycin dose eight times lower than the MIC, underscoring the remarkable efficacy of the combined strategy. This synergy is clinically relevant, as in the absence of antibiotic treatment, the residual bacterial population—although significantly reduced compared to uncoated controls—could still reach levels compatible with infection development in indwelling medical devices. The combination of anti-adhesive coatings with conventional antibiotics therefore provides a promising dual strategy to prevent biofilm formation and device-associated infections.

The coatings also prevented mammalian cell attachment without inducing cytotoxicity, highlighting their dual functionality as anti-adhesive and biocompatible materials. The anti-adhesive properties were attributed to the chemical nature of the amyloid-like aggregates rather than surface roughness or wettability alone. The coatings displayed strong mechanical adhesion and stability under aqueous and mechanical stress, making them suitable for medical devices that require durability during insertion and use. The use of BSA, a readily available and inexpensive protein, combined with a straightforward preparation process, makes this coating strategy economically viable for large-scale medical applications. The results presented in this study suggest a potential strategy to mitigate tube malfunction by reducing both cellular and bacterial adhesion. This approach may prevent blockages caused by cell accumulation and biofilm crystallization, while simultaneously enhancing the effectiveness of conventional antibiotic treatments. Future research could explore the integration of additional antimicrobial agents, such as silver nanoparticles or antibiotics, to further enhance efficacy by local drug delivery.

In conclusion, this work reinforces the potential of BSA amyloid-like coatings as a versatile, biocompatible, and cost-effective strategy to improve the performance and safety of indwelling medical devices, while opening new avenues for multifunctional surface engineering in biomedical applications.

## Acknowledgements

We thank Agencia I+D+I (Projects PICT 2020-2169, PICT Start Up 2020-0034, and PICT 2019-02534), CONICET (Project PIP 0315), and UNLP (Project 11/X948) for their financial support. We are also grateful to Dr. María Esperanza Ruiz for her valuable assistance with the statistical analysis. I.Y.C.G. acknowledges the scholarship granted by CONICET.

## TABLE OF CONTENTS GRAPHIC

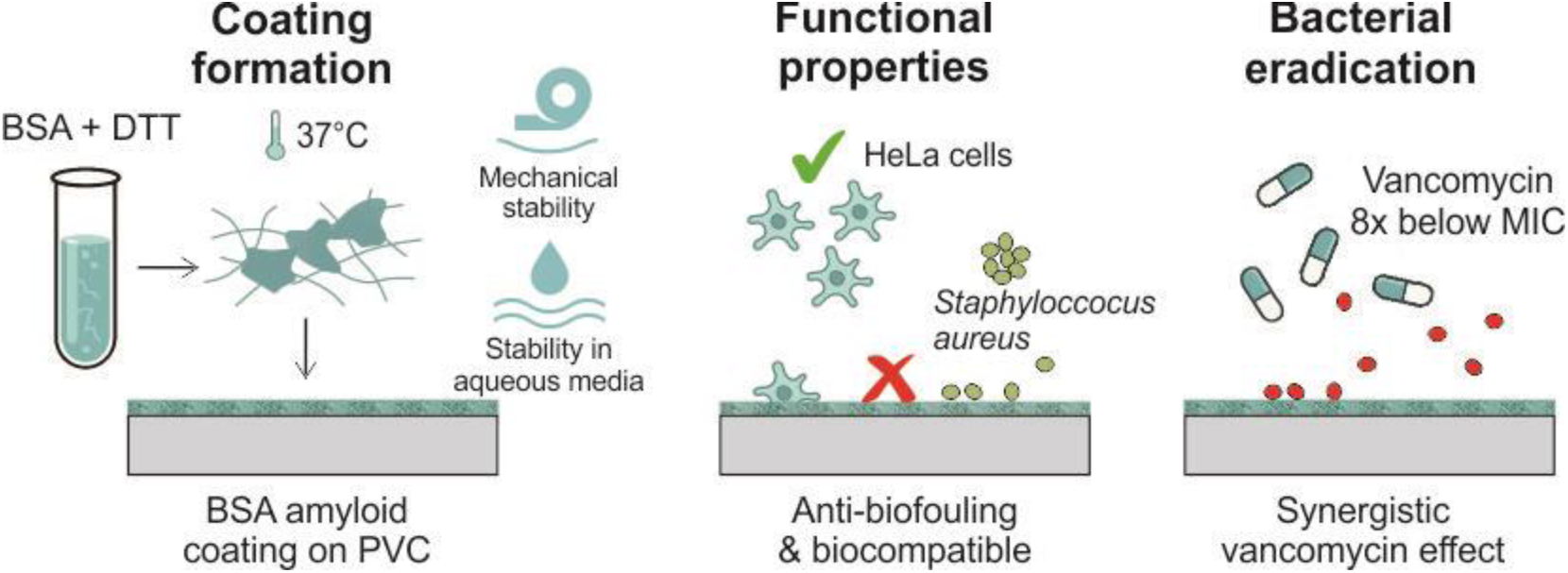

## Notes

### Competing Interest Statement

The authors have declared no competing interest.

## References

(1) *HAIs: Reports and Data* | HAIs | CDC. https://www.cdc.gov/healthcare-associated-infections/php/data/index.html (accessed 2025-01-28).

(2) Bryers, J. D. Medical Biofilms. Biotechnol. Bioeng. 2008, 100 (1), 1–18. 10.1002/BIT.21838.

(3) Ricardo, S. I. C.; Anjos, I. I. L.; Monge, N.; Faustino, C. M. C.; Ribeiro, I. A. C. A Glance at Antimicrobial Strategies to Prevent Catheter-Associated Medical Infections. ACS Infect. Dis. 2020, 6 (12), 3109–3130. 10.1021/ACSINFECDIS.0C00526/ASSET/IMAGES/MEDIUM/ID0C00526_0006.GIF.

(4) Antimicrobial resistance and healthcare-associated infections - Annual epidemiological report 2014 [2012 data]. https://www.ecdc.europa.eu/en/publications-data/antimicrobial-resistance-and-healthcare-associated-infections-annual (accessed 2025-01-28).

(5) Ippolito, M.; Misseri, G.; Catalisano, G.; Marino, C.; Ingoglia, G.; Alessi, M.; Consiglio, E.; Gregoretti, C.; Giarratano, A.; Cortegiani, A. Ventilator-Associated Pneumonia in Patients with COVID-19: A Systematic Review and Meta-Analysis. Antibiot. 2021, Vol. 10, Page 545 2021, 10 (5), 545. 10.3390/ANTIBIOTICS10050545.

(6) Neoh, K. G.; Li, M.; Kang, E. T.; Chiong, E.; Tambyah, P. A. Surface Modification Strategies for Combating Catheter-Related Complications: Recent Advances and Challenges. J. Mater. Chem. B 2017, 5 (11), 2045–2067. 10.1039/C6TB03280J.

(7) Alves, D.; Grainha, T.; Pereira, M. O.; Lopes, S. P. Antimicrobial Materials for Endotracheal Tubes: A Review on the Last Two Decades of Technological Progress. Acta Biomater. 2023, 158, 32–55. 10.1016/J.ACTBIO.2023.01.001.

(8) Kanti, S. P. Y.; Csóka, I.; Jójárt-Laczkovich, O.; Adalbert, L. Recent Advances in Antimicrobial Coatings and Material Modification Strategies for Preventing Urinary Catheter-Associated Complications. Biomed. 2022, Vol. 10, Page 2580 2022, 10 (10), 2580. 10.3390/BIOMEDICINES10102580.

(9) Chen, L.; Song, X.; Xing, F.; Wang, Y.; Wang, Y.; He, Z.; Sun, L. A Review on Antimicrobial Coatings for Biomaterial Implants and Medical Devices. J. Biomed. Nanotechnol. 2020, 16 (6), 789–809. 10.1166/JBN.2020.2942.

(10) Ghilini, F.; Pissinis, D. E.; Miñán, A.; Schilardi, P. L.; Diaz, C. How Functionalized Surfaces Can Inhibit Bacterial Adhesion and Viability. ACS Biomater. Sci. Eng. 2019, 5 (10), 4920–4936. 10.1021/acsbiomaterials.9b00849.

(11) Yang, X.; Guo, J.; Hu, B.; Li, Z.; Wu, M.; Guo, H.; Huang, X.; Liu, X.; Guo, X.; Liu, P.; Chen, Y.; Li, S.; Gu, Y.; Wu, H.; Xuan, K.; Yang, P. Amyloid-Mediated Remineralization in Pit and Fissure for Caries Preventive Therapy. Adv. Healthc. Mater. 2022, 11 (19), 2200872. 10.1002/ADHM.202200872.

(12) Xuan, Q.; Wang, Y.; Chen, C.; Wang, P. Rational Biological Interface Engineering: Amyloidal Supramolecular Microstructure-Inspired Hydrogel. Front. Bioeng. Biotechnol. 2021, 9, 718883. 10.3389/FBIOE.2021.718883/BIBTEX.

(13) Shen, Y.; Levin, A.; Kamada, A.; Toprakcioglu, Z.; Rodriguez-Garcia, M.; Xu, Y.; Knowles, T. P. J. From Protein Building Blocks to Functional Materials. ACS Nano 2021, 15 (4), 5819–5837. 10.1021/ACSNANO.0C08510/ASSET/IMAGES/LARGE/NN0C08510_0008.JPEG.

(14) Kamada, A.; Rodriguez-Garcia, M.; Ruggeri, F. S.; Shen, Y.; Levin, A.; Knowles, T. P. J. Controlled Self-Assembly of Plant Proteins into High-Performance Multifunctional Nanostructured Films. Nat. Commun. 2021 121 2021, 12 (1), 1–10. 10.1038/s41467-021-23813-6.

(15) Levkovich, S. A.; Gazit, E.; Laor Bar-Yosef, D. Two Decades of Studying Functional Amyloids in Microorganisms. Trends Microbiol. 2021, 29 (3), 251–265. 10.1016/J.TIM.2020.09.005.

(16) Balistreri, A.; Goetzler, E.; Chapman, M. Functional Amyloids Are the Rule Rather Than the Exception in Cellular Biology. Microorg. 2020, Vol. 8, Page 1951 2020, 8 (12), 1951. 10.3390/MICROORGANISMS8121951.

(17) Cao, Y.; Adamcik, J.; Diener, M.; Kumita, J. R.; Mezzenga, R. Different Folding States from the Same Protein Sequence Determine Reversible vs Irreversible Amyloid Fate. J. Am. Chem. Soc. 2021, 143 (30), 11473–11481. 10.1021/JACS.1C03392/SUPPL_FILE/JA1C03392_SI_001.PDF.

(18) Krebs, M. R. H.; Devlin, G. L.; Donald, A. M. Amyloid Fibril-like Structure Underlies the Aggregate Structure across the PH Range for β-Lactoglobulin. Biophys. J. 2009, 96 (12), 5013–5019. 10.1016/J.BPJ.2009.03.028/ATTACHMENT/FFF0B979-38F0-4719-891C-06E32BFE3D8E/MMC1.PDF.

(19) Meng, R.; Zhu, H.; Deng, P.; Li, M.; Ji, Q.; He, H.; Jin, L.; Wang, B. Research Progress on Albumin-Based Hydrogels: Properties, Preparation Methods, Types and Its Application for Antitumor-Drug Delivery and Tissue Engineering. Front. Bioeng. Biotechnol. 2023, 11, 1137145. 10.3389/FBIOE.2023.1137145/BIBTEX.

(20) Diaz, C.; Missirlis, D. Amyloid-Based Albumin Hydrogels. Adv. Healthc. Mater. 2023, 12 (7), 2201748. 10.1002/ADHM.202201748.

(21) Arias, Y. A. M.; García, R. R.; Agudelo, M. A. G.; Marín-Pareja, N.; Orozco, C. P. O. Immobilization of Silver Nanoparticles at Varying Concentrations on Segments of Polyvinyl Chloride Manufactured Endotracheal Tubes. Mater. Today Commun. 2024, 41, 110109. 10.1016/J.MTCOMM.2024.110109.

(22) Bernard, L.; Eljezi, T.; Clauson, H.; Lambert, C.; Bouattour, Y.; Chennell, P.; Pereira, B.; Sautou, V. Effects of Flow Rate on the Migration of Different Plasticizers from PVC Infusion Medical Devices. PLoS One 2018, 13 (2), e0192369. 10.1371/JOURNAL.PONE.0192369.

(23) Hu, X.; Tian, J.; Li, C.; Su, H.; Qin, R.; Wang, Y.; Cao, X.; Yang, P. Amyloid-Like Protein Aggregates: A New Class of Bioinspired Materials Merging an Interfacial Anchor with Antifouling. Adv. Mater. 2020, 32 (23), 2000128. 10.1002/ADMA.202000128;WEBSITE:WEBSITE:ADVANCED;CTYPE:STRING:JOURNAL.

(24) Holm, N. K.; Jespersen, S. K.; Thomassen, L. V.; Wolff, T. Y.; Sehgal, P.; Thomsen, L. A.; Christiansen, G.; Andersen, C. B.; Knudsen, A. D.; Otzen, D. E. Aggregation and Fibrillation of Bovine Serum Albumin. Biochim. Biophys. Acta - Proteins Proteomics 2007, 1774 (9), 1128–1138. 10.1016/J.BBAPAP.2007.06.008.

(25) De Vasconcelos, D. N.; Ximenes, V. F. Albumin-Induced Circular Dichroism in Congo Red: Applications for Studies of Amyloid-like Fibril Aggregates and Binding Sites. Spectrochim. Acta Part A Mol. Biomol. Spectrosc. 2015, 150 (1), 321–330. 10.1016/J.SAA.2015.05.089.

(26) Zheng, M.; Pan, M.; Zhang, W.; Lin, H.; Wu, S.; Lu, C.; Tang, S.; Liu, D.; Cai, J. Poly(α-l-Lysine)-Based Nanomaterials for Versatile Biomedical Applications: Current Advances and Perspectives. Bioact. Mater. 2021, 6 (7), 1878–1909. 10.1016/J.BIOACTMAT.2020.12.001.

(27) Kasprzycka, K.; Węglarz, M.; Lewkowicz, M.; Ratuszny, G. The Impact of Quantitative and Semi-Quantitative Culture of Respiratory Tract Secretions on Clinical Decisions in a Patient with Suspected Pneumonia – Case Study. J. Educ. Heal. Sport 2023, 45 (1), 76–85. 10.12775/JEHS.2023.45.01.005.

(28) Fernández-Barat, L.; Motos, A.; Ranzani, O. T.; Bassi, G. L.; Xiol, E. A.; Senussi, T.; Travierso, C.; Chiurazzi, C.; Idone, F.; Muñoz, L.; Vila, J.; Ferrer, M.; Pelosi, P.; Blasi, F.; Antonelli, M.; Torres, A. Diagnostic Value of Endotracheal Aspirates Sonication on Ventilator-Associated Pneumonia Microbiologic Diagnosis. Microorg. 2017, Vol. 5, Page 62 2017, 5 (3), 62. 10.3390/MICROORGANISMS5030062.

(29) Hilt, E. E.; Parnell, L. K.; Wang, D.; Stapleton, A. E.; Lukacz, E. S. Microbial Threshold Guidelines for UTI Diagnosis: A Scoping Systematic Review. Pathol. Lab. Med. Int. 2023, 15, 43–63. 10.2147/PLMI.S409488.

(30) Yang, M.; Dutta, C.; Tiwari, A. Disulfide-Bond Scrambling Promotes Amorphous Aggregates in Lysozyme and Bovine Serum Albumin. J. Phys. Chem. B 2015, 119 (10), 3960–3981. 10.1021/ACS.JPCB.5B00144/SUPPL_FILE/JP5B00144_SI_001.PDF.

(31) LeVine, H. Stopped-Flow Kinetics Reveal Multiple Phases of Thioflavin T Binding to Alzheimer β(1-40) Amyloid Fibrils. Arch. Biochem. Biophys. 1997, 342 (2), 306–316. 10.1006/ABBI.1997.0137.

(32) Arad, E.; Green, H.; Jelinek, R.; Rapaport, H. Revisiting Thioflavin T (ThT) Fluorescence as a Marker of Protein Fibrillation – The Prominent Role of Electrostatic Interactions. J. Colloid Interface Sci. 2020, 573, 87–95. 10.1016/J.JCIS.2020.03.075.

(33) Miller, L. M.; Bourassa, M. W.; Smith, R. J. FTIR Spectroscopic Imaging of Protein Aggregation in Living Cells. Biochim. Biophys. Acta - Biomembr. 2013, 1828 (10), 2339–2346. 10.1016/J.BBAMEM.2013.01.014.

(34) Grigolato, F.; Arosio, P. The Role of Surfaces on Amyloid Formation. Biophys. Chem. 2021, 270, 106533. 10.1016/J.BPC.2020.106533.

(35) Liu, Y.; Tao, F.; Miao, S.; Yang, P. Controlling the Structure and Function of Protein Thin Films through Amyloid-like Aggregation. Acc. Chem. Res. 2021, 54 (15), 3016–3027. 10.1021/ACS.ACCOUNTS.1C00231/ASSET/IMAGES/MEDIUM/AR1C00231_0008.GIF.

(36) Gao, H.; She, J.; Liu, S.; Shi, L.; Lu, X.; Zhang, J.; Wu, C. ‘Green’ Fabrication of PVC UF Membranes with Robust Hydrophilicity and Improved Pore Uniformity. Desalination 2023, 568, 117022. 10.1016/J.DESAL.2023.117022.

(37) Martín, M. L.; Pfaffen, V.; Valenti, L. E.; Giacomelli, C. E. Albumin Biofunctionalization to Minimize the Staphylococcus Aureus Adhesion on Solid Substrates. Colloids Surfaces B Biointerfaces 2018, 167, 156–164. 10.1016/J.COLSURFB.2018.04.006.

(38) Al-Azzam, N.; Alazzam, A. Micropatterning of Cells via Adjusting Surface Wettability Using Plasma Treatment and Graphene Oxide Deposition. PLoS One 2022, 17 (6), e0269914. 10.1371/JOURNAL.PONE.0269914.

(39) Gulati, K.; Adachi, T. Profiling to Probing: Atomic Force Microscopy to Characterize Nano-Engineered Implants. Acta Biomater. 2023, 170, 15–38. 10.1016/J.ACTBIO.2023.08.006.

(40) Hou, Y.; Xie, W.; Yu, L.; Cuellar Camacho, L.; Nie, C.; Zhang, M.; Haag, R.; Wei, Q.; Hou, Y.; Yu, L.; Camacho, L. C.; Nie, C.; Haag, R.; Xie, W.; Zhang, M.; Wei, Q. Surface Roughness Gradients Reveal Topography-Specific Mechanosensitive Responses in Human Mesenchymal Stem Cells. Small 2020, 16 (10), 1905422. 10.1002/SMLL.201905422.

(41) Andrade, J. D.; Hlady, V. Protein Adsorption and Materials Biocompatibility: A Tutorial Review and Suggested Hypotheses. Adv. Polym. Sci. 1986, 1–63. 10.1007/3-540-16422-7_6.

(42) Kikuchi, T.; Matsuura, K.; Shimizu, T. Non-Coating Method for Non-Adherent Cell Culture Using High Molecular Weight Dextran Sulfate and Bovine Serum Albumin. J. Biosci. Bioeng. 2021, 132 (5), 537–542. 10.1016/J.JBIOSC.2021.08.006.

(43) Rha, B.; See, I.; Dunham, L.; Kutty, P. K.; Moccia, L.; Apata, I. W.; Ahern, J.; Jung, S.; Li, R.; Nadle, J.; Petit, S.; Ray, S. M.; Harrison, L. H.; Bernu, C.; Lynfield, R.; Dumyati, G.; Tracy, M.; Schaffner, W.; Ham, D. C.; Magill, S. S.; O’Leary, E. N.; Bell, J.; Srinivasan, A.; McDonald, L. C.; Edwards, J. R.; Novosad, S. Vital Signs: Health Disparities in Hemodialysis-Associated Staphylococcus Aureus Bloodstream Infections — United States, 2017–2020. MMWR. Morb. Mortal. Wkly. Rep. 2023, 72 (6), 153–159. 10.15585/mmwr.mm7206e1.

(44) Ishikawa, K.; Furukawa, K.; Ishikawa, K.; Furukawa, K. Staphylococcus Aureus Bacteremia Due to Central Venous Catheter Infection: A Clinical Comparison of Infections Caused by Methicillin-Resistant and Methicillin-Susceptible Strains. Cureus 2021, 13 (7). 10.7759/CUREUS.16607.

(45) Hurley, J. C. World-Wide Variation in Incidence of Staphylococcus Aureus Associated Ventilator-Associated Pneumonia: A Meta-Regression. Microorg. 2018, Vol. 6, Page 18 2018, 6 (1), 18. 10.3390/MICROORGANISMS6010018.

(46) Kabak, E.; Hudcova, J.; Magyarics, Z.; Stulik, L.; Goggin, M.; Szijártó, V.; Nagy, E.; Stevens, C. The Utility of Endotracheal Aspirate Bacteriology in Identifying Mechanically Ventilated Patients at Risk for Ventilator Associated Pneumonia: A Single-Center Prospective Observational Study. BMC Infect. Dis. 2019, 19 (1), 1–13. 10.1186/S12879-019-4367-7/FIGURES/7.

(47) Bonapace, C. R.; White, R. L.; Friedrich, L. V; Bosso, J. A. Evaluation of Antibiotic Synergy against Acinetobacter Baumannii: A Comparison with Etest, Time-Kill, and Checkerboard Methods. Diagn. Microbiol. Infect. Dis. 2000, 38 (1), 43–50.

(48) Kano, T.; Kawauchi, H. Fibrous Encapsulation of the Peritoneal Catheter in Peritoneal Shunt: Case Report. Surg. Neurol. Int. 2017, 8 (1), 132. 10.4103/SNI.SNI_420_16.

(49) Gaertner, J.; Sabatowski, R.; Elsner, F.; Radbruch, L. Encapsulation of an Intrathecal Catheter. Pain 2003, 103 (1–2), 217–220. 10.1016/S0304-3959(02)00407-4.

(50) Cao, F.; Zhang, L.; Ruan, Y.; Lin, M.; Hong, F. Granuloma Formation after Repeated Episodes of Peritoneal Dialysis Catheter–Related Infection, a Case Report. BMC Nephrol. 2023, 24 (1), 1–3. 10.1186/S12882-023-03230-1/FIGURES/2.

(51) Castañeyra-Ruiz, L.; Lee, S.; Chan, A. Y.; Shah, V.; Romero, B.; Ledbetter, J.; Muhonen, M. Polyvinylpyrrolidone-Coated Catheters Decrease Astrocyte Adhesion and Improve Flow/Pressure Performance in an Invitro Model of Hydrocephalus. Child. 2023, Vol. 10, Page 18 2022, 10 (1), 18. 10.3390/CHILDREN10010018.

(52) Stickler, D. J.; Feneley, R. C. L. The Encrustation and Blockage of Long-Term Indwelling Bladder Catheters: A Way Forward in Prevention and Control. Spinal Cord 2010 4811 2010, 48 (11), 784–790. 10.1038/sc.2010.32.

(53) Yates, A. Using Patency Solutions to Manage Urinary Catheter Blockage. Nurs. Times [online] 2018, 114 (5), 18–21.

